# The neuropeptide Pth2 modulates social behavior and anxiety in zebrafish

**DOI:** 10.1101/2021.07.28.454115

**Authors:** Lukas Anneser, Anja Gemmer, Tim Eilers, Ivan C. Alcantara, Anett-Yvonn Loos, Soojin Ryu, Erin M. Schuman

## Abstract

Animal behavior is strongly context-dependent and behavioral performance is often modulated by internal state. In particular, different social contexts can alter anxiety levels and modulate social behavior. The vertebrate-specific neuropeptide parathyroid hormone 2 (*pth2*) is directly regulated by the presence or absence of conspecifics in zebrafish. As its cognate receptor, the parathyroid hormone 2 receptor (*pth2r*), is widely expressed across the brain, we tested fish lacking the functional Pth2 peptide in several anxiety-related and social paradigms. Rodents lacking PTH2 display increased anxiety-related behavior. Here we show that the propensity to react to sudden stimuli with an escape response is increased in *pth2*^-/-^ zebrafish, consistent with elevated anxiety. While overall social preference for conspecifics is maintained in *pth2^-/-^* fish until the early juvenile stage, we found that both social preference and shoaling are altered later in development. The data presented suggest that the neuropeptide Pth2 modulates several conserved behavioral features, and may thus enable the animal to react appropriately in different social contexts.

## Introduction

Different aspects of animal behavior, such as aggression (Tulogdi *et al*., 2014; Zelikowsky *et al*., 2018; Agrawal *et al*., 2020), anxiety (Meyer *et al*., 2017; Shams *et al*., 2017), or interaction with conspecifics (Shams *et al*., 2018; Groneberg *et al*., 2020), are strongly modulated by social context. In several cases, neuropeptides have been shown to regulate this behavioral plasticity (Zelikowsky *et al*., 2018; Agrawal *et al*., 2020; Gemmer *et al*., 2021). Recently, we found that the expression of the neuropeptide *pth2* (initially described as tuberoinfundibular peptide of 39 residues or *tip39* (Usdin, 1997)) is quantitatively regulated by the density of conspecifics in zebrafish (Anneser *et al*., 2020). In rodents, *pth2* is involved in the regulation of maternal behavior (Coutellier *et al*., 2011), pain processing (Dimitrov *et al*., 2013), oxytocinergic signaling (Cservenák *et al*., 2017), and fear learning (Coutellier and Usdin, 2011). In zebrafish, the cognate receptor of this neuropeptide, *pth2r*, is expressed in approximately 10 % of all neurons, suggesting a potential broad-scale influence (Anneser *et al*., 2020). It has not been described, however, whether the presence or absence of *pth2* alters behavior or other biological features in teleosts.

To identify potential roles of *pth2*, we obtained a line (*pth2^sa23129^*) carrying a premature stop codon which terminates translation at amino acid 55 (out of 157) (Kettleborough *et al*., 2013). In the experiments described below, we tested how the absence of Pth2 influences anxiety-related and social behaviors in larval and late juvenile zebrafish.

## Results

### Pth2^sa23129^ fish lack a functional Pth2 peptide

To investigate the potential roles of Pth2, we in-crossed heterozygous mutants and analyzed the viability of homozygous *pth2^-/-^* fish by observing them for several weeks. We validated the complete absence of Pth2 both by sequencing and by immunostaining (Figure 1). The loss of Pth2 did not lead to differences in either the survival rate or body length (Figure S1). This observation is consistent with data from rodents, in which the deletion of PTH2 did not induce obvious external phenotypic effects except for decreased fertility (Usdin *et al*., 2008). However, in contrast to rodents, matings between *pth2^-/-^* zebrafish led to viable offspring.

**Figure 1.**
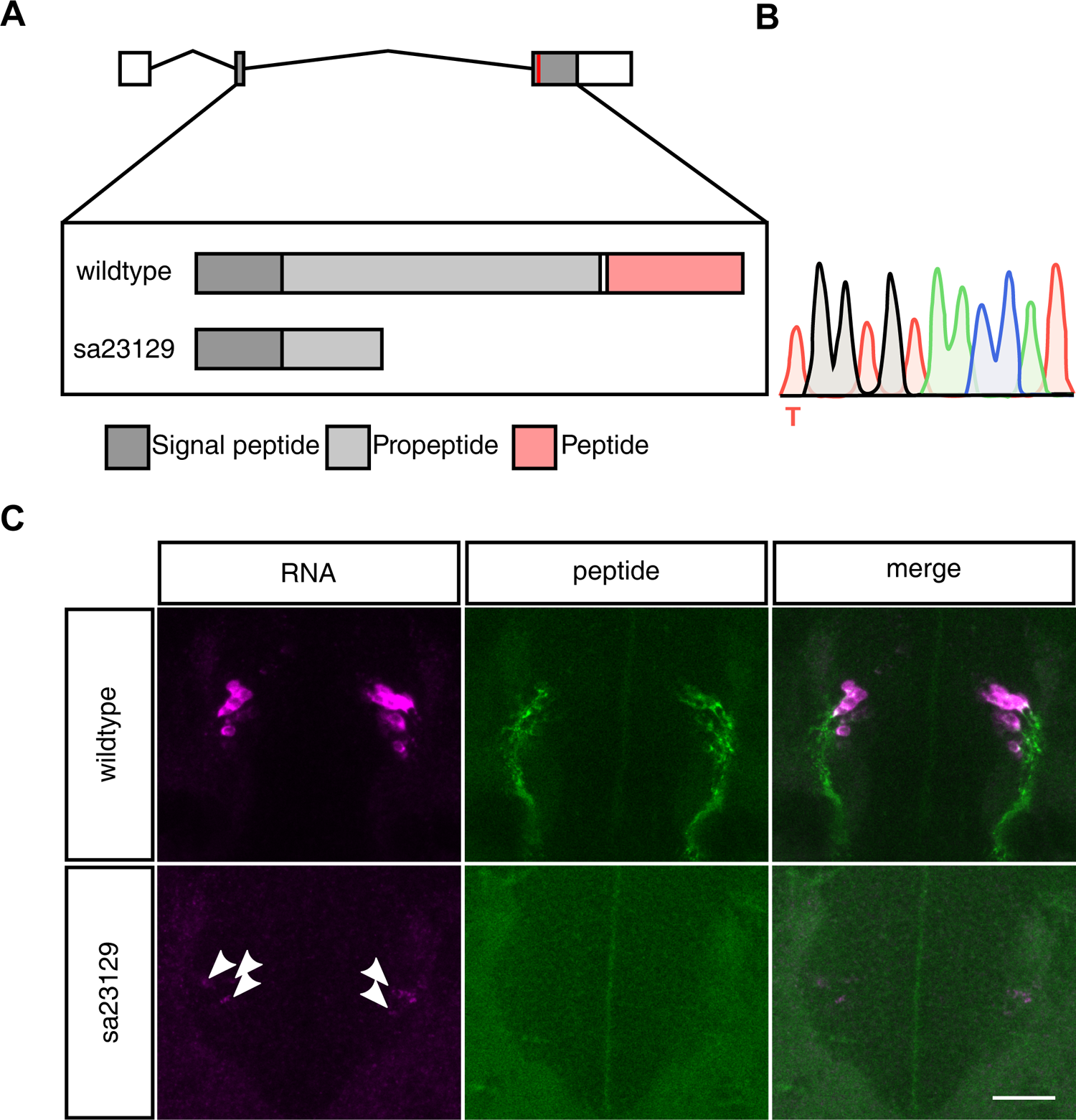
Validation of *pth2*^sa23129^ fish lacking a functional peptide. (A) The *pth2* gene consists of three exons; the coding sequence (CDS) is contained in exon 2 and 3. In *pth2*^sa23129^ fish, a T/A point mutation induces a premature stop codon within the propeptide sequence, thus preventing the successful translation and cleavage of the Pth2 peptide. (B) Homozygous mutants were validated by sequencing and the premature ochre stop codon was identified at base pair 165 out of 474 of the CDS. (C) Transcript levels of *pth2* were strongly decreased in *pth2*^-/-^ animals, as demonstrated by fluorescent in-situ hybridization, also some remaining signal was detected (arrowheads). Abolished Pth2 translation was validated using a custom antibody recognizing a 27-mer epitope immediately preceding the peptide sequence (Anneser *et al*., 2020). For both wildtype and *pth2*^-/-^, 6 larvae at 6 dpf were imaged. Scale bar: 10 µm. See also Figure S1.

### Lack of Pth2 increases startle responsiveness

In rodents, administration of PTH2 has anxiolytic effects (LaBuda, Dobolyi and Usdin, 2004) and animals lacking PTH2 show increased anxiety-related behavior (Fegley *et al*., 2008). A common assay to investigate anxiety-related behavior in zebrafish is the startle response (Burgess and Granato, 2008; Reider and Connaughton, 2015; Tomasi *et al*., 2020), in which fish are presented with a stimulus and react with stereotyped escape responses, so-called C-starts (Burgess and Granato, 2007). Previous work showed that higher rearing densities in zebrafish decreased the likelihood of a startle response (Burgess and Granato, 2008). As *pth2* levels are strongly influenced by the number of conspecifics (Anneser *et al*., 2020), we tested *pth2^-/-^* and *pth2^+/+^* fish at 5 days post fertilization (dpf) in a startle paradigm and measured when and how often they responded to a vibrational cue with an escape (Figure S2A and Figure 2A). We observed a typical bimodal distribution characterizing the startle response with fish either exhibiting a short-or long-latency response (Burgess and Granato, 2007) (SLC or LLC, Figure 2B) mediated by Mauthner-cells (Burgess and Granato, 2007) or a prepontine cell population, respectively (Marquart *et al*., 2019). When we compared the time-to-response after stimulus onset between *pth2^-/-^* and *pth2^+/+^* animals, we found SLCs occurred moderately but significantly earlier in mutants (Figure 2C), suggesting that *pth2^-/-^* fish respond more quickly to potential threats (Troconis *et al*., 2017). In general, *pth2^-/-^* fish showed a higher overall startle response rate (see Figure 2D). We analyzed how the proportion of SLCs, LLCs, and non-responding animals per trial were related to the genotype of the fish with an ANOVA and found significant interactions between these ratios, indicating that for both SLCs and LLCs, *pth2*^-/-^ fish showed higher response rates (Figure 2E). Furthermore, in a logistic regression model relating response ratios and genotype we found the increased overall response rate of mutant fish to be a significant predictor of the *pth2^-/-^* genotype (Figure S2B-D).

**Figure 2.**
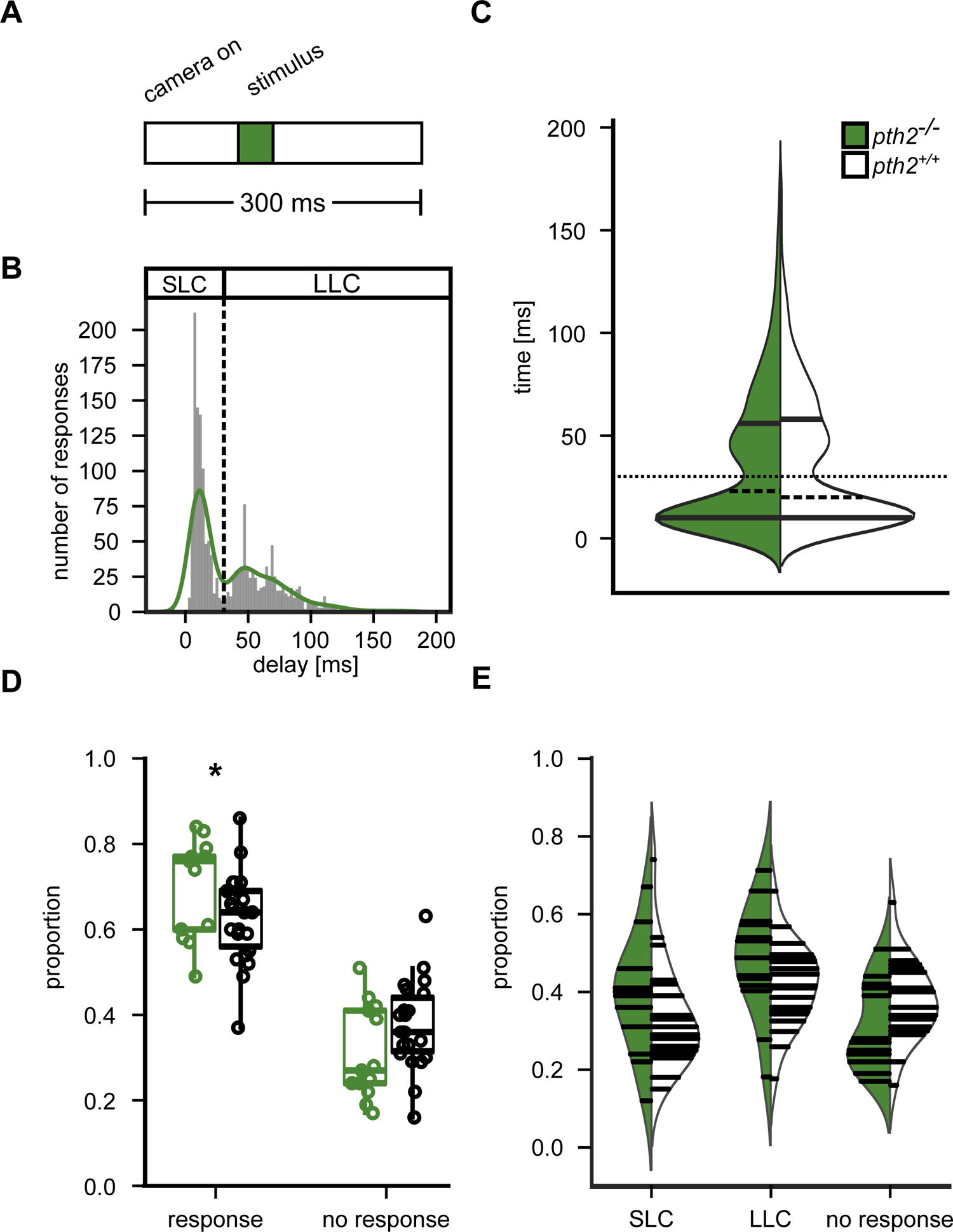
*pth2*^-/-^ fish display increased startle propensity. (A) Experimental scheme. Ten larval fish (5 dpf) were recorded with 500 Hz for 300 ms. After a baseline of 100 ms, a 40 ms pulse was delivered using a bass shaker. (B) The histogram and overlaid kernel density estimate show the delay with which animals respond after the onset of the stimulus with a C-start. We recorded 1613 escape responses from 220 wildtype fish and observed the bimodal distribution characteristic of Mauthner-cell-mediated short-latency C-starts (SLC)(Burgess and Granato, 2007) and long-latency C-starts (LLC) mediated by a prepontine cell group(Marquart *et al*., 2019). (C) Delay after stimulus until escape onset is shown as a violin plot for wildtype and *pth2*^-/-^ animals. Median is shown as dashed lines and the quartiles as solid lines. The overlaid dotted line indicates the boundary between SLCs and LLCs. For *pth2*^-/-^ fish, 1330 startle responses were recorded from 130 larvae. Both genotypes displayed a similar bimodal distribution, however, a Kolmogorov-Smirnov test indicated a moderate yet significant difference (p_D=0.082_= 0.0001). We then tested SLCs and LLCs separately with a Mann-Whitney-U test and found SLCs in *pth2^-/-^* fish to occur earlier (p_U=253197.0_= 9.17e-09) and LLCs to be indistinguishable between genotypes (p_U=235143.5_= 0.11). (D) Box plots show the fraction of animals that react to a stimulus with an escape response. Fractions were calculated for groups of 10 animals, with n = 13 for *pth2*^-/-^ and n = 22 for wildtype. The proportion of *pth2*^-/-^ animals performing an escape response as reaction to a startle stimulus was significantly increased as compared to wildtype (Mann-Whitney-U test, p_t=92.0_ = 0.042). (E) The violin plot shows the proportion of *pth2*^-/-^ and wildtype animals responding to a startle stimulus with an SLC, LLC, or do not react at all. As these proportions are directly dependent upon each other, we found the interaction between all response types to be significantly related to the genotype in a logistic regression model (p_z=2.01_ = 0.044). See also Figure S2.

### Maintenance of social preference is disrupted in *pth2^-/-^* fish

As *pth2* levels are strongly influenced by the social environment of an animal (Anneser *et al*., 2020), we queried whether the absence of *pth2* might alter social preference for conspecifics or other features of social interaction. We first tested the propensity of animals to stay close to conspecifics in two different paradigms. First, we placed animals in a U-shaped chamber which allowed the fish to freely explore both arms (Dreosti *et al*., 2015); at the end of one arm the fish could see conspecifics, separated by a transparent wall. After a habituation period, three conspecifics were placed in one of the arms and the time the experimental fish spent in the corresponding area was measured (Dreosti *et al*., 2015) (see Figure 3A). At the early juvenile stage (21 dpf), both *pth2^+/+^* and *pth2^-/-^* animals showed a clear preference for the arm containing the conspecifics and no difference between genotypes was apparent (Figure 3B). As the development and maintenance of social preference is modulated by social context well after 21 dpf (Gemmer *et al*., 2021), we raised wild-type and mutant animals until the late juvenile stage (56 dpf) and tested them again. At this stage, while *pth2^+/+^* animals still displayed a strong preference for the conspecific-containing arm, *pth2^-/-^* fish did not (Figure 3C). Consistent with these results, we found that rearing of wild-type fish in isolation (which reduces *pth2* levels) abolished the social preference as well (Figure 3D). The second social paradigm we tested used a rectangular tank, in which the animals could see three age-matched fish, placed in one of two adjacent areas, from any location. We used the fraction of time the animals spent in the half of the arena closer to the social area to measure social preference (Figure 3E). Just as we observed in the U-shaped chamber, no difference was found at 21 dpf, but lack of *pth2* led to a trend for a social preference decrease at 56 dpf, although it did not reach the significance threshold (Figure 3F, G). Likewise, social isolation of wild-type fish recapitulated these effects (Figure 3H).

**Figure 3.**
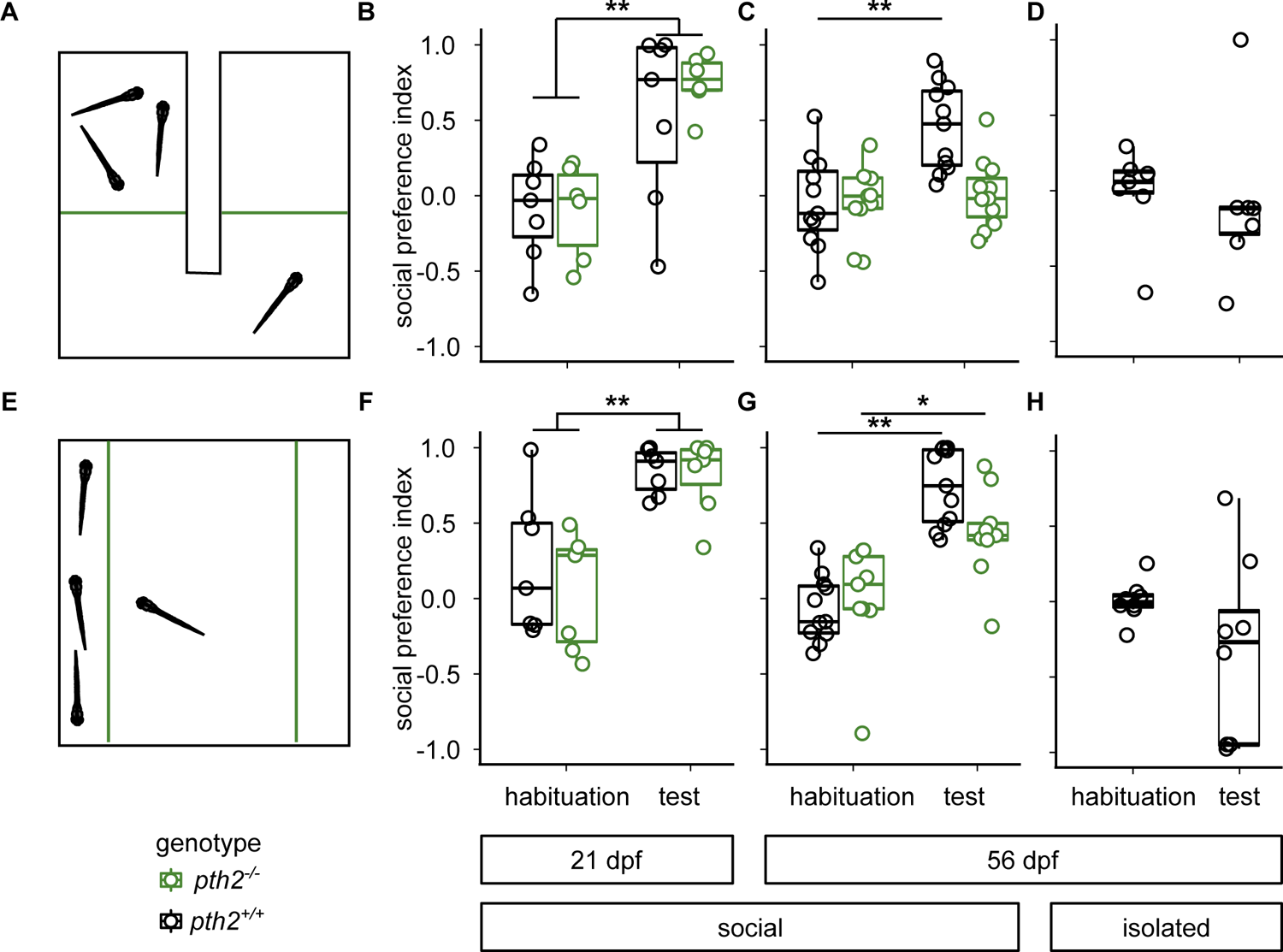
Maintenance of social preference is disrupted in *pth2*^-/-^ mutants. (A) Scheme of the U-shaped chamber, in which 21 dpf fish could spend time in either of two adjacent compartments, separated by acrylic glass (dotted line)(Dreosti *et al*., 2015). Visual access to three age-matched wildtype conspecifics was only possible in one of the two areas. (B) Box plots indicate the preference for the social area displayed by *pth2^+/+^* (n = 7) and *pth2*^-/-^ fish (n = 6) at 21 dpf during habituation and in response to the actual presence of stimulus fish. While animals showed no preference during habituation, they spent a significantly increased amount of time close to conspecifics during the test period (two-way ANOVA, p_F=22.42_ = 0.0001). The genotype of the animals was not found to influence behavior (two-way ANOVA, p_F=0.46 =_ 0.505). (C) At 56 dpf, both the presence of conspecifics (two-way ANOVA, p_F=11.39_= 0.002) and the genotype of the animals (two-way ANOVA, p_F=7.21_= 0.01; n = 11 for both genotypes) contributed significantly to social preference. Additionally, we found a significant interaction between these factors (p_F=7.79_= 0.008). Therefore, we performed post-hoc Tukey-Kramer tests and found that *pth2^+/+^* fish displayed a significant increase in SPI under test conditions (p_q=6.17_= 0.001), whereas *pth2^-/-^* fish did not (p_q=0.58_= 0.9). (D) Box plots show the effect of rearing animals in isolation on social preference in the forced-choice paradigm. In 56 dpf animals, the presence of conspecifics led to no obvious change in behavior (n = 7; paired, one-sided t-test: p_t=0.58_= 0.58). (E) Scheme for open field social preference test, in which fish could move freely in a rectangular dish, whilst having visual access to three conspecifics. (F) Box plots indicate social preference of wildtype (n = 7) and *pth2*^-/-^ fish (n = 7) at 21 dpf during habituation and in response to stimulus fish. The presence of conspecifics during the test period altered behavior of all animals significantly (two-way ANOVA, p_F=30.949_ = 0.00001), while genotype did not influence behavior (two-way ANOVA, p_F=0.268_ = 0.609). (G)At 56 dpf, the presence of conspecifics (p_F=48.76_=3.45e-08) and its interaction with the test animal’s genotype (p_F=4.49_=4.09e-02) significantly altered social preference (for *pth2^-/-^*, n = 9, for *pth2^+/+^*, n = 8). In a post-hoc Tukey-Kramer test, we found *pth2*^+/+^ fish to strongly prefer the arm with conspecifics as compared to habituation (p_q=9.34_= 0.001). In *pth2*^-/-^ fish the same effect was present, although less pronounced (p_q=4.4_= 0.018). (H) After 8 weeks of social isolation, *pth2^+/+^* fish displayed no social preference towards conspecifics (one-sided, paired t-test: p_t=1.38_= 0.21, n = 8).

### Shoal cohesion is decreased in *pth2^-/-^* fish

Zebrafish groups display collective behavior, characterized by coordinated swimming of individuals (Miller and Gerlai, 2012). Several individual genes have been shown to contribute to this behavior (Tang *et al*., 2020). As our previous results indicated that *pth2* is regulated by the presence of conspecifics, we tested whether its absence might influence collective behavior. At 56 dpf, groups of 20 *pth2^+/+^* or *pth2^-/-^* fish were placed in a circular tank (diameter of 70 cm) and their behavior was recorded for 30 minutes. Animals of either genotype immediately started to adapt their movement to the group and explored the entire tank together (Figure 4A, Figure S3A). Across the duration of the trials, pth2^+/+^ fish reached higher average velocities (Figure 4B). As zebrafish move faster in more polarized groups (Couzin *et al*., 2002; Miller and Gerlai, 2012; Tang *et al*., 2020), we analyzed cohesiveness and alignment of the animals. We analyzed inter-individual distances including the nearest neighbor distance, the median distance and the maximum distance, and found that the maximum distance between individuals was consistently increased in *pth2^-/-^* animals (Figure 4C, Figure S3B), suggesting a lesser degree of cohesion. To measure the alignment between the animals, we computed the degree to which individual trajectories could be explained by the movement of the group centroid (Figure 4D). To this end, we computed a polarization parameter *p* that was determined by calculating the variance of the matrix consisting of all 20 individual trajectories explained by the two first principal components obtained in a principal component analysis, which closely reflected centroid movement. This polarization parameter was significantly lower in *pth2^-/-^* animals across all trials (Figure 4E-F), further supporting the notion that collective motion of *pth2^-/-^* fish occurs in a less coordinated way. When we counted the number of times that individual animals clearly broke away from the main group, we also found that both the total number of excursions and the time spent away from the main group were higher in groups of *pth2^-/-^* fish (see Figure 4G, H). Ultimately, we used several features of group motion in a dimensionality reduction approach and found that *pth2^-/-^* and *pth2^+/+^* replicates preferentially occupy different regions in the transformed space (see Figure 4I, Figure S3D, E). This separation of genotypes was mainly driven by features related to the spacing of the fish, such as the average distance between animals and the area occupied by the entire group (see Figure 4J, Figure S3C, D).

**Figure 4.**
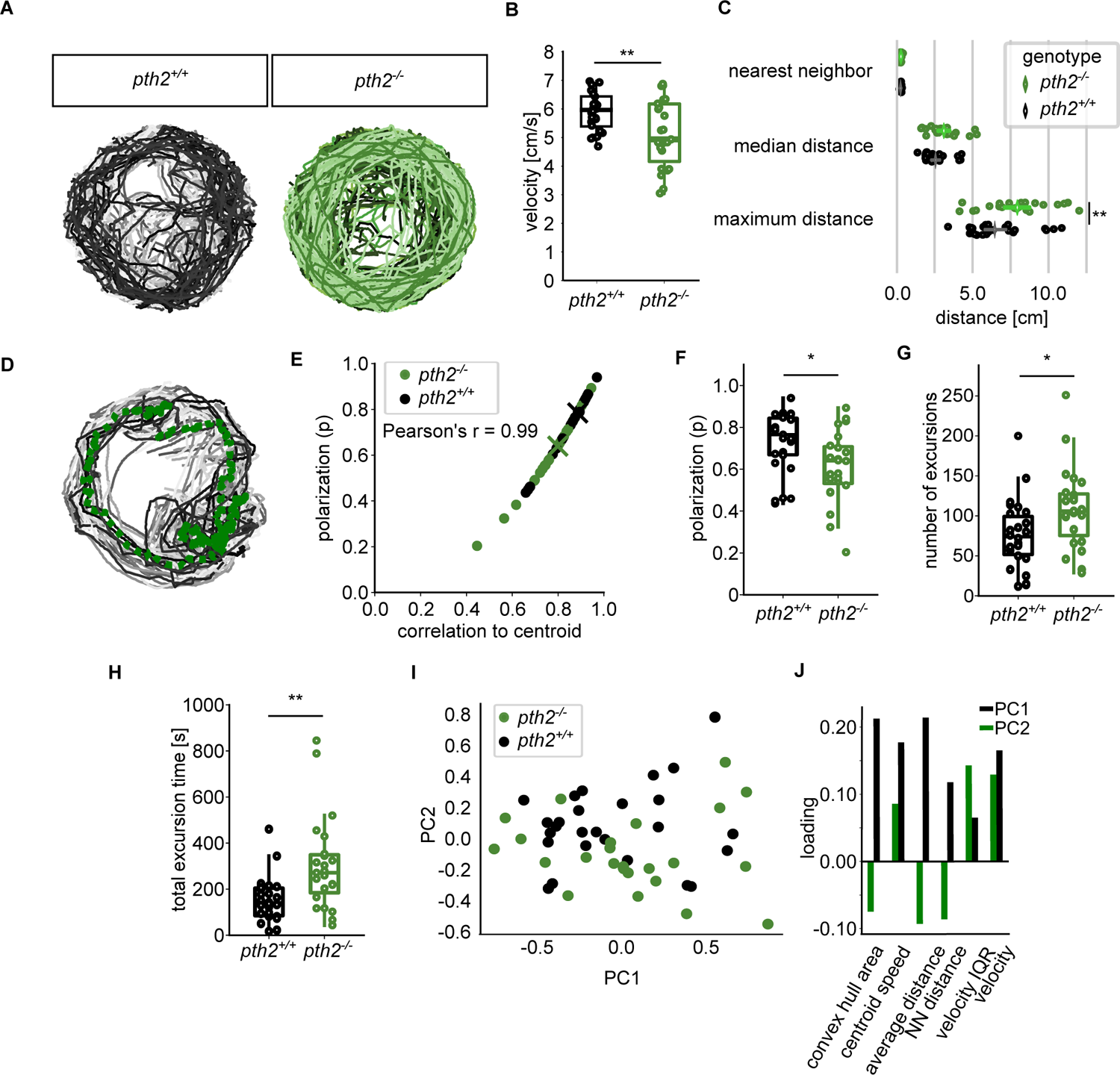
Fish move in a less cohesive manner in groups of *pth2^-/-^* animals. (A) Example traces of 20 fish over one minute in the behavioral tank. Fish movement was strongly coordinated across all replicates. For *pth2*^-/-^, n = 26, for wildtype, n = 29. (B) Box plots depict the median movement speed of fish over 30 minutes. Wildtype animals moved on average faster than fish without *pth2* (unpaired, one-sided t-test, p_t=2.938_ = 0.005). (C) Dot plots highlight the increased distance between *pth2*^-/-^ fish when compared to wildtype animals. In an ANOVA, genotype was found to be a significant factor for distance (p_F=9.85_=0.002), Tukey’s post-hoc range test then showed the maximum distance was significantly different between genotypes (p_q=5.21_=0.004). (D) Trajectories of wildtype animals over one minute with the trajectory of the shoal centroid overlaid as green dotted line. (E) Polarization of animals was strongly correlated with the mean of the individual animals’ correlation to the shoal centroid. Median values for *pth2^-/-^* and *pth2^+/+^* are indicated by the colored x (Pearson’s r = 0.99). (F) The animals’ polarization was significantly increased in wildtype animals (unpaired, one-sided t-test, p_t=2.62_= 0.011). (G)Wildtype animals were more likely to consistently move in coherent groups, as evidenced by the increased propensity of *pth2^-/-^* animals to split away from the main group (unpaired, one-sided t-test, p_t=2.51_= 0.016). (H) The smaller number of excursions in wildtype animals was not compensated by a longer total excursion time (unpaired, one-sided t-test, p_t=3.40_= 0.001). (I) Projecting several features into the space of the first two principal components after dimensionality reduction led to a preferred location of *pth2^+/+^* replicates in the upper half of PC space. (J) The loadings of the principal components used in (k) indicate that the separation of *pth2^+/+^* and *pth2^-/-^* is mainly driven by distance-related features, such as the convex hull area, average distance, and nearest-neighbor distance (NN). See also Figure S3.

## Discussion

In many systems, social context modulates the abundance of neuropeptides and thus influences physiology and behavior (Pan *et al*., 2009a; Slavich and Cole, 2013; French *et al*., 2016; Zelikoswky, Ding and Anderson, 2018). Here we show that the socially regulated neuropeptide *pth2* influences anxiety-related and social behaviors in zebrafish. *pth2* has been shown to regulate similar behaviors in other species, suggesting an evolutionarily conserved role for this peptide across vertebrates. For example, several lines of evidence suggest the involvement of *pth2* in the regulation of anxiety: intracerebroventricular administration of PTH2 in rodents led to an increase of time spent in the open arms of an elevated plus maze (LaBuda, Dobolyi and Usdin, 2004). In other experiments, mice lacking either PTH2 or its receptor showed elevated levels of freezing when placed in a context where they received foot shocks before (Coutellier and Usdin, 2011). We showed that zebrafish lacking Pth2 have a higher propensity to react to sudden stimuli with an escape response. Startle responses are stereotyped, but can be modulated and states of elevated anxiety have been shown to increase startle responsiveness in rats (Walker and Davis, 1997) and humans (Grillon *et al*., 1999). Interestingly, social isolation led to the same phenomenon (Wilkinson *et al*., 1994; Parr, Winslow and Davis, 2002), suggesting *pth2* to be involved in the regulation of anxiety states.

Increased anxiety has often been associated with social isolation. A common finding is that rodents tend to visit the open arms in elevated plus mazes less after experiencing social isolation (Pan *et al*., 2009b; Kumari *et al*., 2016; Nakagawa *et al*., 2019). The acquisition and retention of fear memory has been shown to be enhanced in socially-isolated animals (Lukkes *et al*., 2009; JH *et al*., 2015; Zelikowsky *et al*., 2018). As protection from predators by diluting the risk of being targeted (Hamilton, 1971; Vine, 1971) is one of the key benefits of sociality it is a logical consequence of social isolation to increase vigilance and respond to potentially threatening stimuli at a lower threshold (Hawkley and Cacioppo, 2010). For example, fruit flies display a graded decrease in freezing in response to threatening stimuli that is proportional to group size (Ferreira and Moita, 2020). Our results suggest that Pth2 might contribute to the regulation of appropriate vigilance states in vertebrates.

In three different paradigms assessing social interaction, we found *pth2^-/-^* animals to display altered behavior. In zebrafish social preference develops with age and reaches peak levels at around three weeks of development (Dreosti *et al*., 2015). Here we showed, that up until 21 dpf, no difference was found between *pth2^-/-^* and *pth2^+/+^* animals in either of two social preference paradigms. However, 56 dpf *pth2^-/-^* animals show a clear decrease in social preference, suggesting a role for *pth2* in the maintenance of social preference at later developmental stages. Previous studies have shown that social preference can be altered by rearing conditions (Tunbak *et al*., 2020; Gemmer *et al*., 2021), which also affect the maintenance of social preference at later developmental stages (Gemmer *et al*., 2021). Consistent with these findings, *pth2^-/-^* animals form less cohesive shoals than *pth2^+/+^* fish and swim in a less polarized manner. Features of collective behavior such as group alignment and shoal cohesion are genetically controlled (Tang *et al*., 2020; Aspiras *et al*., 2021), suggesting that *pth2* might be one of the effector genes that contributes to the regulation of group behavior. So far, the impact of *pth2* on behavior has only been analyzed in depth in rodents. Social behavior in mammals is quite distinct from fish, so that a direct comparison between previous results and ours is hard to draw. Previous studies have convincingly shown that Pth2 contributes to maternal behavior (Cservenák *et al*., 2010, 2013; Gellén *et al*., 2017; Dobolyi, Cservenák and Young, 2018) and the posterior intralaminar thalamus, where one of the thalamic Pth2+ populations is found in mammals, integrates social cues, eventually activating oxytocinergic neurons (Cservenák *et al*., 2017; Dobolyi, Cservenák and Young, 2018; Valtcheva *et al*., 2021).

Neuropeptidergic modulation of neuronal circuits is an ancient motif of neuronal information processing (Jékely, 2013, 2021) and although adaptation of neuropeptidergic circuits for new functions is common, some neuropeptides are remarkably consistent in the kind of modulation they induce (Minakata, 2010). Both zebrafish and human Pth2 peptides efficiently activate Pth2r in the other species, arguing for a well-conserved system (Papasani *et al*., 2004). We provide evidence that lack of Pth2 induces changes in anxiety levels and social behavior in zebrafish, behaviors that have been shown to be affected by Pth2 in other species. In several other paradigms, social behavior and anxiety have been found to be co-modulated (Felix-Ortiz *et al*., 2016; Zou *et al*., 2016; Zelikowsky *et al*., 2018). Pth2 might thus serve a conserved function in the regulation of these important behavioral domains.

### Limitations of the Study

In this study, we explored the effects of a loss-of-function mutation in the gene *pth2* on the startle response and social behavior in zebrafish. Although we observed clear behavioral changes in startle responsiveness, we did not test whether improved sensory perception or a reduced threshold in the effector circuits is responsible for the behavioral difference. Additionally, the startle response is just one of many possible measures of anxiety levels in zebrafish. This study does not test whether the absence of Pth2 leads to similar effects in other behavioral settings. In our experiments testing social preference, we found that the absence of Pth2 leads to a decline in social preference in 56 dpf old animals, which we could recapitulate by raising animals in isolation. However, we did not investigate whether at this stage short-term isolation (which also shuts down Pth2 transcription) is sufficient for disrupting social preference. Finally, in our shoaling experiment we did not investigate individual differences due to uncertainties about tracking individual fish for the entirety of the trial. Further work will be necessary to elucidate the precise forms of interaction deficits in *pth2^-/-^* fish.

## Methods

### Animal Stock and Husbandry

Adult zebrafish of the lines Konstanz wildtype (KN) and Pth2^sa23129^ (RRID:ZIRC_ZL12243.19, referred to as *pth2*^-/-^) were kept at 28°C on a light cycle of 14-hour light/10-hour dark and housed in 3.5 L ZebTEC tanks at a density of 5-35 fish of mixed sexes. Heterozygous Pth2^sa23129^ fish were obtained from the European Zebrafish Resource Center (EZRC, Karlsruhe, Germany) and outcrossed to our standard wildtype strain three times to reduce potential background mutations. Fish were then maintained as homozygous mutants. Fish were fed with brine shrimp *(Artemia salina)* and/or GEMMA Micro three times per day. In addition, vinegar eelworms (*Turbatrix aceti*) were fed to larval and juvenile fish. Larvae up to 5 days post fertilization (dpf) were kept in dishes filled with E3 medium (5 mM NaCl, 17 mM KCl, 0.33 mM CaCl2, 0.33 mM MgSO4) in a 28°C incubator also in a 14-hour light/10-hour dark cycle. All animal procedures conformed to the institutional guidelines of the Max Planck Society and were approved by the Regierungspräsidium Darmstadt, Germany (governmental ID: V 54-19 c 20/15-F126/1016).

### Genotyping

To validate the genotype of the animals used in our experiments, all fish were genotyped after completion of the experiments. Genomic DNA was extracted from parts of the tail by placing tissue in 50 µl 50 mM NaOH. Samples were heated to 95 °C for 15 minutes under vigorous shaking. Tubes were cooled to 4 °C and 5 µl of 1 M Tris-HCl (pH 8.0) were added for neutralization. Tubes were centrifuged for 5 minutes at 8,000 rcf and 1 µl supernatant was used for PCR. The pth2^sa23129^ fish are characterized by a T→A point mutation in the *pth2* gene that leads to a premature stop codon at amino acid 55/157. The flanking region was amplified using the primer pair 5’-GCTGTTAGGCGAGTGTC-3’ and 5’-CTATGTTTCTCTTCTGCTGGTGAC–3’, genotype was then determined by sequencing.

### Whole-mount in-situ hybridization and immunohistochemistry

Fixation and staining were performed as described previously (Herget *et al*., 2014). Pth2 mRNA was visualized using a previously published in-situ probe (Anneser *et al*., 2020). A custom antibody raised in guinea pig against zebrafish Pth2 (Anneser *et al*., 2020) was used in a dilution of 1:500. Secondary antibodies were tagged with Alexa594 dyes and used in a dilution of 1:1000. After in-situ hybridization and immunohistochemistry, animals were transferred stepwise into 87% glycerol and mounted dorsally for imaging using an inverted confocal microscope (LSM-780, Zeiss, Jena, Gemany). For all conditions, animals were imaged with a 20x air objective, using a 488 nm (92 µW) or a 594 nm laser (39 µW).

### Startle Response

For this behavioral paradigm, animals were reared in specified densities of 10 animals per 20 ml of E3 in dishes of 6 cm diameter as of 3 dpf. At 5 dpf, fish were transferred within their dish to the behavioral chamber, which was placed in a fully enclosed steel casing that reduced outside visual and auditory stimulation. After 5 minutes of habituation, the experiment commenced. Vibrational cues were delivered with a Monacor EX-1W bass shaker (Conrad Electronics, 1594623), which was activated by a function generator (Hewlett Packard, 3312A Function Generator), amplified by a Kemo M034N power amplifier (Conrad Electronics, 191010). Stimulus frequency was set to 70 Hz and generated as a sine wave with an intensity pre-determined to induce startle responses in approximately 70 % of wildtype fish. Delivery was controlled by an Arduino Mega 2560 Rev3 (Arduino), which jointly triggered the bass shaker and a Redlake MotionXtra HG-SE high-speed camera, which recorded at 500 Hz at 512×512 pixel resolution. Homogeneous illumination was achieved with an LED-72T ring light around the objective, placed 10 cm above the setup. 100 ms after camera onset, a 40 ms startle cue was delivered. In total, animal behavior was recorded for 300 ms. This was repeated 10 times with a 15 second inter-trial interval, which was previously reported to prevent habituation (Burgess and Granato, 2007). Using a custom-written MATLAB script, we manually assessed the onset of startle-responses, which were defined by the characteristic C-shape in which the animal bends(Burgess and Granato, 2007). Startle responses in zebrafish occur in two waves, the short-latency C-start (SLC) and the long-latency C-start (LLC). Using the latency histogram of the escape onset, we found the typical bimodal distribution reported previously (Burgess and Granato, 2007) and determined the trough between the two kinds of responses to occur at 30 ms after stimulus onset. This value was then used to distinguish between SLCs and LLCs. The average fraction of fish responding with an SLC, an LLC, or not at all was computed across all ten trials for each experiment.

### Social Preference Testing

Fish were raised in densities of 30 fish per 3.5 liter until 21 dpf. At this day, animals were placed in one of two behavioral chambers. For the open field social preference task, the setup consisted of a rectangular chamber (height 10 mm) with a test area of 25 x 75 mm and adjacent stimulus areas of 8 x 75 mm (21 dpf) or 50 x 75 mm and 16 x 75 mm (56 dpf). For the other social preference task, animals were placed in a U-shaped chamber of previously published design (Dreosti *et al*., 2015), with a total area of 40 x 32 mm. Areas for the stimulus fish were 15 x 15 mm in dimension and the width of the connection between the two arms was 6 mm (21 dpf). At 56 dpf, total area was 72 x 60 mm with stimulation area of 28 x 28 mm and the width of the connection 15 mm. Water level was adjusted to 5 mm. White backlight illumination was provided from below with a computer screen and the animals were recorded at a temporal resolution of 20 Hz. To reduce outside stimulation, behavioral chambers were placed in a sound-absorbing steel box. Before each experiment, the chambers were cleaned with hot water (∼ 60°C) and refilled with fresh ZebTEC stand-alone system water (28.5°C). Experimental fish were placed in the middle of the behavioral chamber and recorded for 10 minutes using the Pylon recorder software without any stimulus fish present (habituation period). Afterwards, three stimulus fish were placed in one of the adjacent stimulus areas and behavior was recorded for another 15 minutes. To compute the social preference index, the location of the animal was tracked using custom-written software and the fraction of time the animals spent either in the “social” half of the test area (open field social preference) or in the arm of the U-shaped chamber which allowed visual access to conspecifics was calculated in MATLAB.

### Collective Behavior

Fish were raised until 2 months of age at densities of 50 fish per 3.5 L. To observe collective behavior, 20 fish were placed in a setup consisting of a white round exploration tank with a diameter of 70 cm. The tank was placed in a basin filled with 29.5°C water (heated by a pump) to maintain the temperature of the water inside the behavioral chamber for the 30 minutes of experiment. A high-resolution camera (Basler acA4112-30um), positioned 73 cm above the chamber, recorded the fish (30 frames per second) using the Pylon recorder software (https://software.scic.brain.mpg.de/projects/PylonRecorder/PylonRecorder) and 1200 white LEDs in the ceiling provided homogeneous illumination (approximately 650 Lux). A white box surrounding the setup reduced visual and acoustic disturbances of the fish. For each replicate, the chamber was cleaned and refilled with 3.0 liter 28.5°C warm system water. Before the experiments, home tanks were moved from the ZebTEC system to a 28.5°C incubator in the experimental room. Approximately 10 minutes before the experiment started, 20 fish of similar size were transferred from their home tank to a 1L breeding cage (TECNIPLAST) with a nursery insert (TECNIPLAST, Part Number: ZB300BTI). This way, all 20 fish were moved to the center of the behavioral chamber at once. Recording started immediately and continued for 30 minutes. Individual trajectories were obtained using the TRex software(Walter and Couzin, 2021) with a tracking threshold of 80.

### Outcome measures

For the collective behavior experiment, several measurements were computed from the obtained individual trajectories. For each frame, all pairwise distances between the 20 fish were computed. From this distance matrix, we computed the median of the nearest-neighbor (NN), the median of the maximum-distance and the median distance across all fish. Velocity was calculated as the distance travelled between consecutive frames, and then recorded as cm per second. Ultimately, the median velocity across all fish was computed. Additionally, the inter-quartile range for velocity was calculated to obtain a measurement for consistency of movement. Cumulative shoal distance was calculated by computing the centroid of the group of fish and cumulatively adding the distance this centroid covered between consecutive frames. To analyze group cohesion, we calculated the area occupied by the smallest convex hull we could fit around the group using standard algorithms (Barber *et al*., 1996) implemented in scipy (Virtanen *et al*., 2020). Additionally, we calculated dispersion of the animals as follows: we first calculated the randomized nearest-neighbor distance by temporally shuffling locations of animals within a video (NN_shuffled_). Dispersion was then computed according to the following formula: dispersion = log(NN / NN_shuffled_) (Harpaz *et al*., 2021). To further examine the coordination of swimming, we calculated a polarization parameter *p* as follows: X- and y-components of the individual trajectories were assembled in a 40 x 54000 matrix and decomposed to obtain the first two principal components (PCs), which largely resembled the trajectory described by the centroid of the group. The sum of the variance explained by these two PCs was calculated to obtain the parameter *p*, which allowed us to exactly quantify the degree to which the individual trajectories were correlated. Across the videos, we often observed individual animals or small groups that split away from the main group. To automatically identify these animals, we modified an approach described before by Miller and Gerlai in 2011 (Miller and Gerlai, 2011). In short, we calculated for each video the overall distribution of nearest-neighbor distances (dNND) and used the mode of the distribution as a first threshold to extract movement segments for each individual, during which their nearest-neighbor distance exceeded this threshold. For each segment, the maximum nearest-neighbor distance was stored and used to compute a second, maximum nearest-neighbor distribution. We used the p = .015 quantile of this distribution as a threshold to automatically identify proper excursions. To account for different sizes of splinter groups, we replaced *d*NND with NND_n_ – NND_n-1_, with n being the splinter group size. For example, if two fish moved away from the group, their respective NND would be small, but the distance to the second closest neighbor large. Since we recorded 20 fish, we obtained 10 different excursion types, which we then binarized for further analysis. We used this algorithm to compute the number of excursions performed during one experiment, the total time the animals spent away from the main group, the average excursion time and the average number of fish taking part in an excursion. To evaluate the extent to which the animals display thigmotaxis, we computed a boldness score, for which we assessed what fraction of the total time the animals spent in the middle of the arena. We computed two concentric circles, the outer one being the boundary of the arena, the inner one being defined by a radius which was the outer circle’s radius divided by the square root of 2. This implied that the area covered by the inner circle was equal to the area of the outer circle minus the inner circle’s area and if the animals had no preference for any part of the arena, they should spent 50 % of their time in each of the circles.

## Data Analysis

Significance values are reported as follows: * p < 0.05, ** p < 0.01 and *** p < 0.001. All replicates are biological replicates obtained from different breeding pairs. For the social preference tasks, we performed ANOVA to analyze whether the variables genotype of the animals and presence of conspecifics significantly altered the social preference index. In case of significant interactions, Tukey-Kramer post-hoc tests were performed. For the startle response, we compared the average fraction of animals per experiment that responded to stimulation with an escape response with the fraction of non-responders using an unpaired, one-sided t-test. We then segmented the responses into SLCs and LLCs and fit a logistic regression model relating genotype with the individual response variables using a subset of the data. We bootstrapped the accuracy of the model by repeated sampling and evaluated with a hold-out test-set containing 30% of the entire dataset. To visualize the model, we projected the logistic decision boundary we obtained with the entire dataset onto the first two principal components of the Eigen-reduced matrix containing SLC, LLC, and no-response rates for all experimental replicates.

The collective behavior data was analyzed as follows: Cumulative shoal distance, velocity, boldness, dispersion, the polarity parameter p and hull area were compared between genotypes using one-sided, unpaired t-tests. To test nearest-neighbor distance, median distance, and maximum distance between animals, we first conducted an ANOVA to see whether the genotype affected these features. We then used the post-hoc Tukey-Kramer test to identify the features affected by genotype. Following previous approaches (Tang *et al*., 2020), we combined several features (velocity, nearest-neighbor distance (NN), median distance, convex hull area, velocity interquartile-range, and centroid speed) into a matrix and scaled all features to the range between 0 and 1 (Pedregosa *et al*., 2011). We visualized this matrix as a heatmap using the Euclidean distance between replicates for hierarchical clustering. To further determine whether this range of features could help to distinguish between wildtype and mutants, we used dimensionality reduction methods and plotted the points in the space determined by the two first principal components of a PCA and the first two dimensions we obtained from a t-SNE reduction. Unpaired, one-sided t-tests were used to compare all other values.

The survival of animals was analyzed by comparing the Kaplan-Meier Estimates of *pth2*^-/-^ and *pth2*^+/+^ fish animals using a log-rank test (Kaplan and Meier, 1958), their size at 14 dpf was measured from snout to the onset of the tail fin and compared with a one-sided, unpaired t-test.

All data analysis was performed using custom-written python (python 3.7) scripts in the jupyter notebook 6.0.0 environment, embedded in the anaconda navigator 1.9.7 (64-bit version). Graphs and figures were compiled in Inkscape 0.92. Results are reported as box plots with individual data points overlaid, as violin plots either with quantiles indicated or with individual data points on top, or as scatter plots with the mean and its confidence interval (0.95) overlaid.

## Acknowledgements

Florian Vollrath wrote the MATLAB script to track startle responses and compute social preference. Norman Heller and Andreas Umminger built the chamber for analysis of startle behavior, Fabian Bayer, Erik Papuschin, and Dr. Claudio Polisseni built the tracking setup for collective behavior. Friedrich Kretschmer discussed important aspects of data analysis with us.

## Author Contributions

L.A. and E.M.S. conceived the project. L.A., A.G., T.E., and I.C.A. performed the experiments. L.A., A.G. and T.E. analyzed the data. A.-Y. L. and S.R. provided valuable input throughout the study. L.A. and E.M.S. wrote the manuscript and prepared the figures. All authors read and reviewed the manuscript.

## Data Availability

All data is openly available from the EDMOND repository https://dx.doi.org/10.17617/3.6v

## Code Availability

All relevant analysis code is available from the GitHub repository https://github.com/Anneser/zfish_PTH2

---

**Figure S1.**
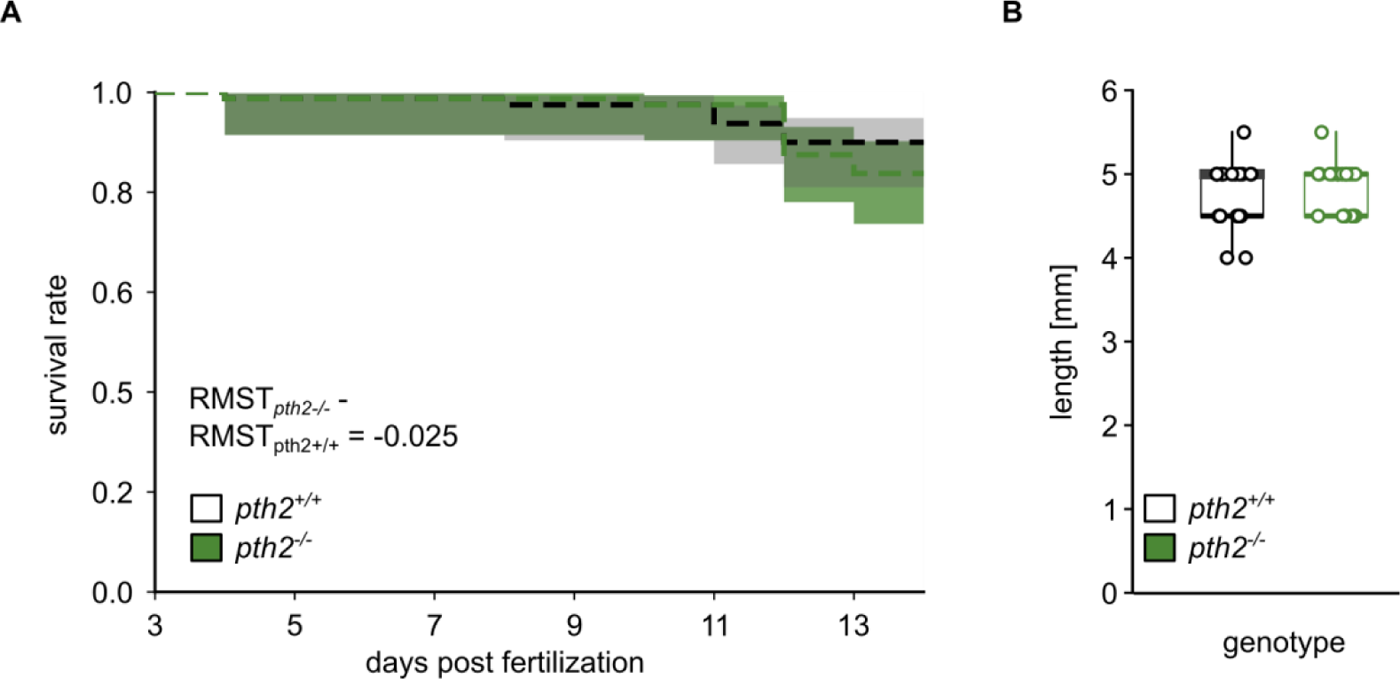
Survival rate and growth are not affected by *pth2.* Relates to Figure 1. (A) Kaplan-Meier curve shows the survival rate of *pth2^-/-^* and wildtype fish over the first two weeks of development. Shaded areas indicate the .95 confidence interval. For both groups, n = 80. We analyzed the curves with a log-rank test and found no significant difference (p_χ2=1.15_= 0.28), supported by a difference in mean restricted survival times between groups of 0.025 days. (B) Box plots highlight the average body length at 2 weeks post fertilization. For both groups, n = 20. Distributions were compared using a one-sided, unpaired t-test (p_t=0.7_= 0.48).

**Figure S2.**
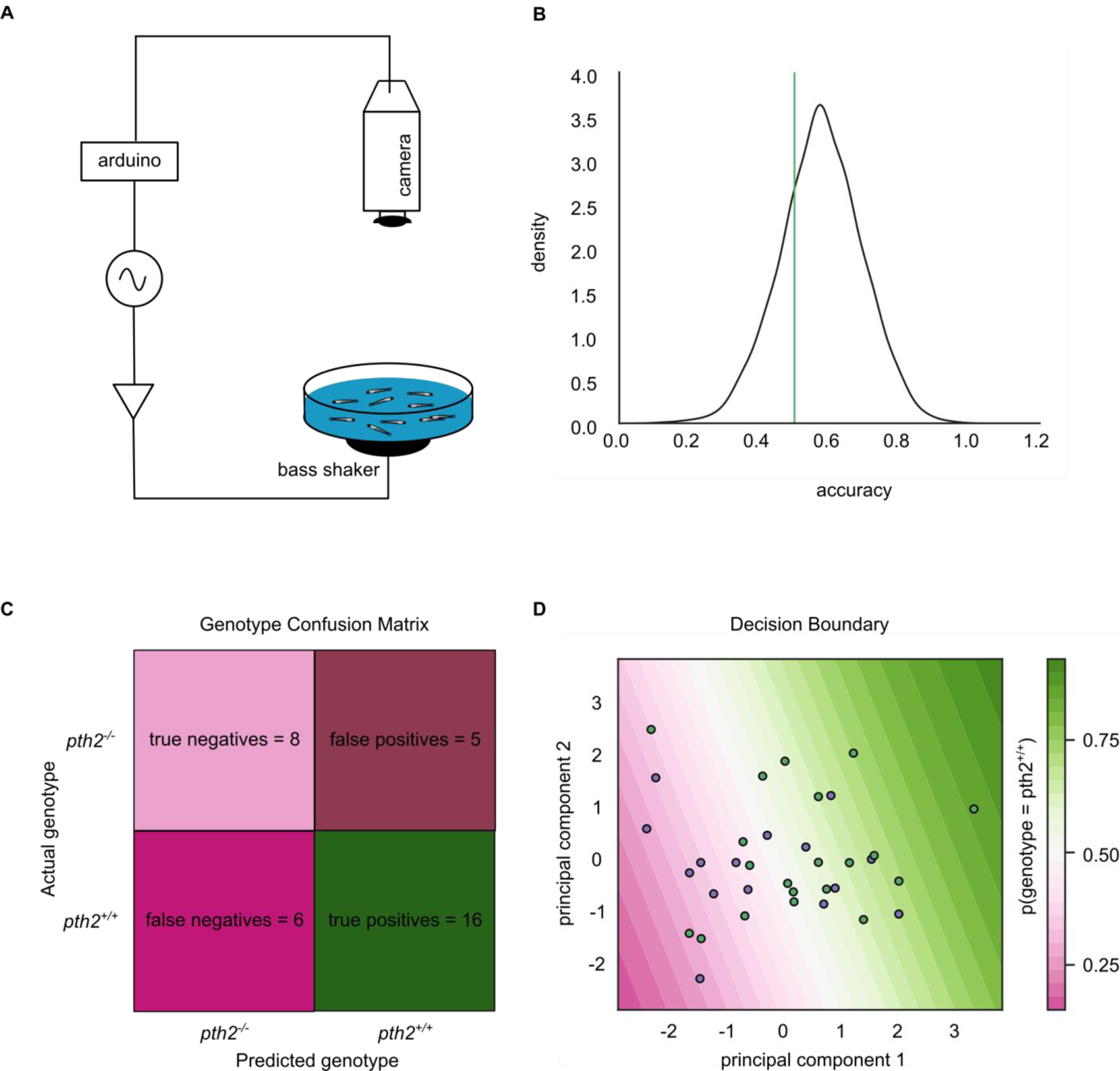
In-depth analysis of startle response. Relates to Figure 2. (A) Experimental scheme for the startle experiment. Both the bass shaker for startle cue delivery and the high-speed camera were triggered by an Arduino board at the same time to ensure alignment of videos and cue. (B) Graph shows the bootstrapped accuracy of the logistic regression model fit to the data. We trained the model 1,000 times sampling 70 % of the full dataset as training set and testing the model on the remaining 30 % as test set. The vertical green line indicates chance level of correct assignment of genotype. (C) Confusion matrix of the full model, highlighting the number of true and false predictions. (D) The fraction of SLCs, LLCs, and failures to respond were projected onto the space of the first two principal components to facilitate visualization. Wildtype are plotted as green dots, *pth2^-/-^* in magenta. The background color indicates the decision boundary computed by the logistic regression model.

**Figure S3.**
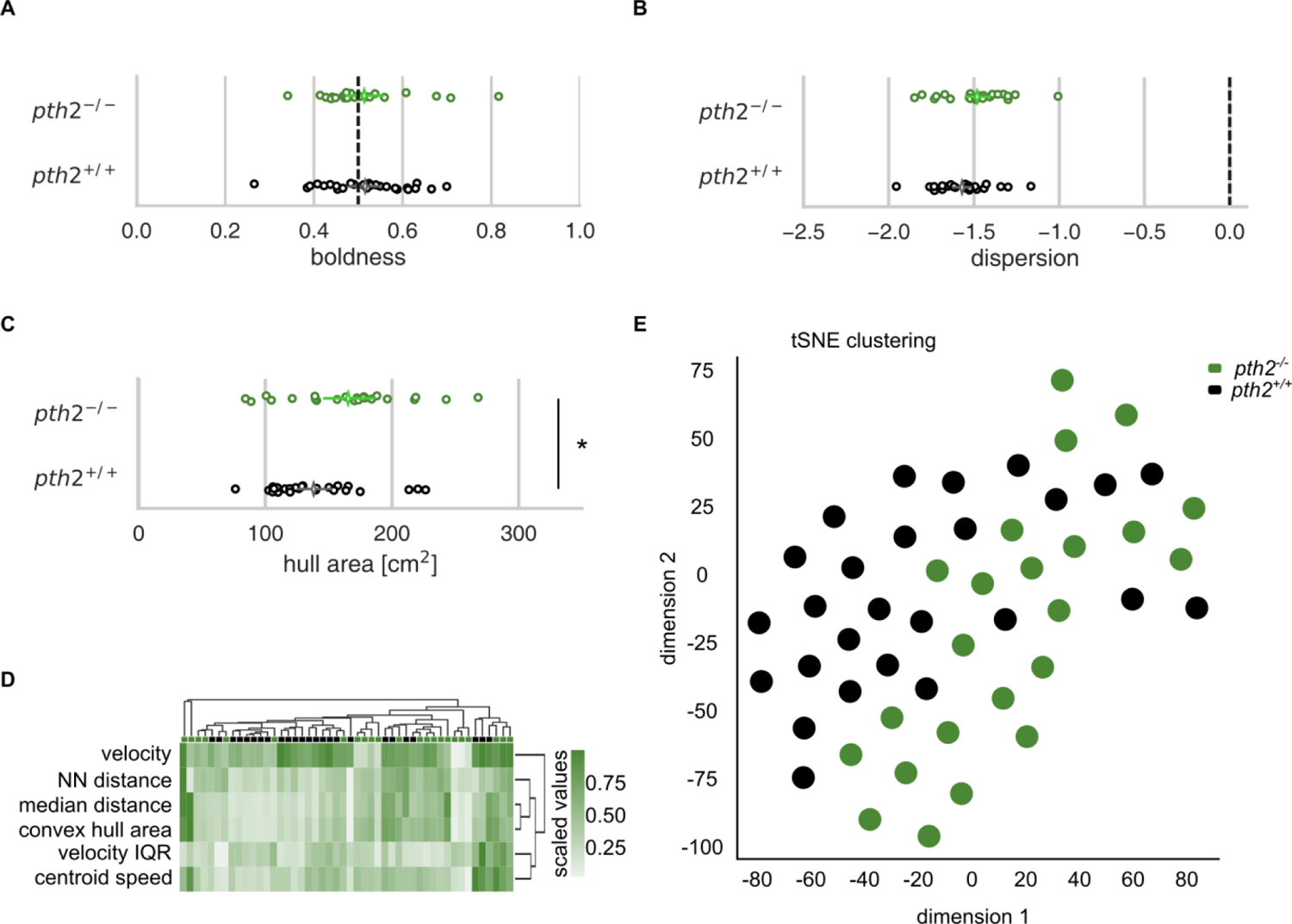
In-depth analysis of shoaling behavior. Relates to Figure 4. (A) The boldness parameter indicates the fraction of time the animals spend in the center of the arena versus the periphery. Chance level is indicated by the dashed vertical line. No difference was found between groups (one-sided, unpaired t-test, p_t=0.07_=0.94). (B) Dispersion relates the median nearest-neighbor distance to a random distribution (random distance indicated by dashed vertical line)(Harpaz *et al*., 2021). No difference was detected between groups (one-sided, unpaired t-test, p_t=1.74_= 0.08). (C) Dot plot shows the median area occupied by a convex hull covering all 20 fish over 30 minutes as a measure of shoal cohesion. The mean and its .95 confidence interval are overlaid. For all graphs in this figure, n = 26 for *pth2^-/-^* and n = 29 for *pth2^+/+^*. Groups were compared with a one-sided, unpaired t-test (p_t=2.24_= 0.03). (D) Heatmap shows the scaled values of several features of fish behavior across individual replicates (*pth2^+/+^* depicted in black, *pth2^-/-^* in green). Feature matrix was hierarchically clustered and re-ordered based on the obtained dendrogram. Scatter plot shows the matrix displayed in (DS) after t-distributed stochastic neighbor embedding. While *pth2^+/+^* replicates preferentially occupy the left part of the space, *pth2^-/-^* are separated along dimension 1, recapitulating the effect demonstrated in Figure (4)I.

